# A quantitative evaluation of MIRU-VNTR typing against whole-genome sequencing for identifying *Mycobacterium tuberculosis* transmission: A prospective observational cohort study

**DOI:** 10.1101/252734

**Authors:** David Wyllie, Jennifer Davidson, Tim Walker, Preeti Rathod, Derrick Crook, Tim Peto, Esther Robinson, Grace Smith, Colin Campbell

## Abstract

**Background:** Mycobacterial Interspersed Repetitive Unit-Variable Number Tandem Repeat (MIRU-VNTR) typing is widely used in high-income countries for *Mycobacterium tuberculosis* typing. Whole-genome sequencing (WGS) is known to deliver greater specificity, but no quantitative prospective comparison has yet been undertaken.

**Methods:** We studied isolates from the English Midlands, sampled consecutively between 1 January 2012 and 31 December 2015. In addition to routinely performed MIRU-VNTR typing, DNA was extracted from liquid cultures and sequenced using Illumina technology. Demographic and epidemiological data were extracted from the Enhanced Tuberculosis Surveillance system maintained by Public Health England. Closely related samples, defined using a threshold of five single nucleotide variants (SNVs), were compared to samples with identical MIRU-VNTR profiles, with shared epidemiological risk factors, and to those with both characteristics.

**Findings:** 1,999 patients were identified for whom at least one *M. tuberculosis* isolate had been MIRU-VNTR typed and sequenced. Comparing epidemiological risk factors with close genetic relatedness, only coresidence had a positive predictive value of over 5%. Excluding co-resident individuals, 18.6% of patients with identical MIRU-VNTR profiles were within 5 SNVs. Where patients also shared social risk factors and ethnic group, this rose to 48%. Only 8% of MIRU-VNTR linked pairs in lineage 1 were within 5 SNV, compared to 31% in lineage 4.

**Interpretation:** In the setting studied, MIRU-VNTR typing and epidemiological risk factors are poorly predictive of close genomic relatedness, assessed by SNV. MIRU-VNTR performance varies markedly by lineage.

**Funding:** Public Health England, National Institute of Health Research Oxford Biomedical Research Centre.

## Research in context

### Evidence before this study

We searched Pubmed using the search terms ‘whole genome sequencing’ and ‘MIRU-VNTR’ and ‘tuberculosis’ for English language articles published up to December 21^st^, 2017. Multiple studies have shown that most pairwise genomic comparisons will be within five SNVs when direct transmission has occurred from one individual to another. Both outbreak studies and population studies have demonstrated how whole-genome sequencing generates smaller clusters than MIRU-VNTR typing, and how sequence data allows for differentiation of isolates within a cluster. However, no systematic comparison of MIRU-VNTR typing vs. WGS has however been published. The degree to which WGS provides more specific results, and the degree to which it is likely to be more cost effective, therefore remains uncertain.

### Added value of this study

This study seeks to quantify the predictive value of identical MIRU-VNTR profiles, and of overlapping demographic and epidemiological data, for close genomic relatedness in a cosmopolitan setting. Importantly, it demonstrates that in our setting MIRU-VNTR-based clustering predicts genomic relatedness differently depending on *M. tuberculosis* lineage. Whether this is due to biological differences between the lineages or to immigration patterns, it is likely that these findings are relevant to other cosmopolitan settings. These data provide an explanation as to why MIRU-VNTR typing was not cost-effective when implemented in England, and indicate WGS may perform substantially better.

### Implications of all the available evidence

Whilst it is generally accepted that WGS provides more informative results than MIRU-VNTR typing, the latter is still practiced widely under the belief that it remains a helpful tool for public health investigations. This study shows that whilst differing MIRU-VNTR profiles help exclude close genomic relatedness, matching profiles rarely predict such relatedness. Having quantified its predictive value at a population level, this study should hasten the transition from MIRU-VNTR typing to WGS in other settings similar to ours.

## Introduction

In 2016 there were 5,664 notified cases of tuberculosis in the England, with an incidence of 10.2 per 100,000 population.^1^ Despite a steady fall in incidence since its peak early this decade, this remains the highest rate in western Europe, outside of the Iberian peninsula.^2^ Much of the recent decline in incidence has been due to a falling number of patients born outside of the UK. However, this decline slowed in 2016, with domestic transmission likely to still be contributing towards the residual case load.

Rapid detection of *Mycobacterium tuberculosis* transmission should offer enhanced opportunities for disease control.^3,4^ In England, as in many high-income countries, tuberculosis transmission has been identified with the help of Mycobacterial Interspersed Repetitive Unit-Variable Number Tandem Repeat (MIRU-VNTR) typing, which clusters cultured isolates on the basis of their molecular fingerprints.^5,6^ A recent post-deployment evaluation of the MIRU-VNTR-based surveillance programme in England has however questioned the cost-effectiveness of this approach.^7^

Since 2015, Public Health England has been undertaking a phased introduction of routine whole genome sequencing (WGS) for all mycobacterial cultures.^8^ This has meant the relatedness of isolates could be simultaneously compared using both single nucleotide variants (SNV) and by MIRU-VNTR typing, and has provided a novel opportunity to compare the added value of whole genome sequencing (^9-14^;Table 1) in an unselected population, at scale.

**Table 1.** Previous studies including both MIRU-VNTR and SNV analysis of M. tuberculosis

| Samples | Comment | Reference |
| --- | --- | --- |
| 36 archived Manila strain isolates | SNV analysis revealed variation not demonstrated by MIRU-VNTR. | <sup>9</sup> |
| 390 retrospective isolates from the English Midlands | Genetic heterogeneity within MIRU-VNTR clusters demonstrated. 5 and 12 SNV proposed as potential cut offs for epidemiological relatedness. | <sup>10</sup> |
| 199 epidemiologically linked cases sequenced retrospectively | Relationship with MIRU-VNTR profile was not addressed | <sup>22</sup> |
| 36 isolates from an outbreak | SNV analysis revealed variation not demonstrated by MIRU-VNTR. | <sup>23</sup> |
| 50 cases from an outbreak | SNV analysis revealed variation not demonstrated by MIRU-VNTR. | <sup>11</sup> |
| 1,000 isolate sample of 2,248. Representative of Russian population studied, plus 28 diverse sequences | Relationship with MIRU-VNTR profile was not addressed. Multiple sub-lineages observed within Lineage 4 (Euro-American). | <sup>24</sup> |
| 69 cases from an outbreak defined by a SNV | SNV analysis revealed variation not demonstrated by MIRU-VNTR. | <sup>12</sup> |
| 86 cases from an outbreak | SNV analysis revealed variation not demonstrated by MIRU-VNTR. | <sup>13</sup> |
| 90 cases belonging to 35 MIRU-VNTR clusters | MIRU-VNTR performance overestimated transmission particularly in immigrants infected with closely related strains | <sup>14</sup> |

Here we estimate what proportion of *M. tuberculosis* isolates from a cosmopolitan area of central England that are linked by MIRU-VNTR typing, or have associated epidemiological risk factors, are closely genomically related.

## Methods

### Samples studied for comparison of MIRU-VNTR with SNVs

Consecutive *M. tuberculosis* isolates from the Public Health England Centre for Regional Mycobacteriology Laboratory, Birmingham between 1 January 2012 and 31 December 2015 were included in the study. This laboratory serves a large catchment of approximately 12 million persons in the English Midlands, a region which includes high, medium (40-150 cases per 100,000 population), and low TB incidence areas.

### Identification and MIRU-VNTR typing

Clinical samples were grown in Mycobacterial Growth Indicator tubes (MGIT) (Becton Dickinson, New Jersey, USA), and *M. tuberculosis* was identified using Ziehl-Neelsen staining, followed by nucleic acid amplification and hybridisation using Genotype Mycobacterium CM hybridisation tests (Hain LifeScience, Nehren, Germany). MIRU-VNTR typing^5^ was performed on the first isolate from each patient in each calendar year, following protocols then in place.

### Laboratory and bioinformatic processing

This was carried out as described.^10^ Nucleic acid was extracted from 1•7 ml of MGIT culture as described.^8^ Illumina 150 bp paired end DNA libraries were made using Nextera XT version 2 chemistry kits and sequenced on MiSeq instruments (Illumina). Reads were mapped to the H37Rv v2 reference genome (Genbank: NC000962.2) using Stampy^15^, and aligned to Bam files parsed with Samtools mPileup^16^, with further filtering performed based on the base and alignment quality (q30 and Q30 cutoffs, respectively). SNV variation was reported but indels were not considered as part of this work as they have been reported to be less reliably called than SNVs.^15^ Bases supported only by low confidence base calls were recorded as uncertain (‘N’), as were positions with > 10% minor variant frequencies, and all calls at the genomic positions included in Supplementary Data 1, since these regions were repetitive (as identified by self-self blastn analysis) or were found to commonly contain low-confidence mapping (*rrl*, *rrs*, *rpoC* and *Rv2082* loci). Such uncertain bases were ignored in pairwise SNV computations.

### Metrics of relatedness

We used pairwise SNV distances between isolates as a metric of close genetic relatedness, considering isolates closely genetically related when their pairwise SNV distance was less a particular SNV threshold. For the main analysis, 5 SNV was used as the threshold, but a range of other thresholds were considered in sensitivity analyses.

Lineage assignation was performed using ancestral SNVs, as described.^17^ Relatedness between samples was determined by comparing the number of mismatching positions between loci using BugMat.^18^ Relatedness between MIRU-VNTR profiles compared the total number of differences in repeat lengths at each of the 24 loci. For example, for a one-locus typing scheme, if isolate 1 had 3 repeats, and isolate 2 had 5 repeats, we coded this as a 2 MIRU-VNTR repeat unit difference.

### Collection and collation of patient data

Demographic data (sex, age, ethnic group and residence), and social risk factor data (current or history of imprisonment, drug misuse, alcohol misuse or homelessness) were obtained from the Enhanced Tuberculosis Surveillance system. Co-residence was defined as having the same first line of address and postcode.

### Statistical analyses

We considered a series of categorical variables as predictors of close genomic relatedness in logistic regression analyses. Additionally, for some variables, we constructed composite categorical variables reflecting whether more than one risk factor was present. For each given SNV threshold, we estimated odds ratios for close genomic relatedness using logistic regression. Separately, we modelled the relationship between SNV variation (s) (outcome), *Mycobacterium tuberculosis* lineage (l, a discrete variable) and n, the number of MIRU-VNTR repeat number differences observed, as defined above. We modelled

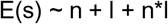

thus allowing estimation of both lineage-specific variation in the absence of any variation in MIRU-VNTR types, and how SNV increased with increasing MIRU-VNTR differences. We used quantile regression (R quantreg package) for the main analysis as homoscedascity assumptions were violated. All analyses used R 3.3.1 for Windows.

### Ethical framework

Public health action taken as a result of notification and surveillance is one of the Public Health England‘s key roles as stated in the Health and Social Care Act 2012 and subsequent Government directives which provide the mandate and legislative basis to undertake necessary follow-up. Part of this follow-up is identification of epidemiological and molecular links between cases.This work is part of service development carried out under this framework, and as such explicit ethical approval is unnecessary.

### Funding source

This study is supported by the Health Innovation Challenge Fund (a parallel funding partnership between the Wellcome Trust [WT098615/Z/12/Z] and the Department of Health [grant HICF-T5-358]) and NIHR Oxford Biomedical Research Centre. Professor Derrick Crook is affiliated to the National Institute for Health Research Health Protection Research Unit (NIHR HPRU) in Healthcare Associated Infections and Antimicrobial Resistance at University of Oxford in partnership with Public Health England. Professor Crook is based at University of Oxford. The views expressed are those of the author(s) and not necessarily those of the NHS, the NIHR, the Department of Health or Public Health England. The sponsors of the study had no role in study design, data collection, data analysis, data interpretation, or writing of the report. The corresponding author had full access to all the data in the study and had final responsibility for the decision to submit for publication.

## Results

### Isolates studied

We studied all *M. tuberculosis* isolates consecutively grown in, or referred to, the Public Health England Mycobacterial reference centre for the English Midlands between 2012-2015 (n=2,718) (Figure 1). We excluded 551 isolates because MIRU-VNTR typing had already been performed on another isolate (protocol was to MIRU-VNTR type one isolate per patient per year), and 57 isolates because of technical concerns about laboratory processing (Figure 1). The remaining 2,110 isolates came from 2,020 discrete patients. A further 16 isolates were excluded because multiple isolates from the same individual were separated by >12 single nucleotide variants (SNVs) (suggestive of technical error), along with five recurrent cases of *M. tuberculosis* infection, leaving 1,999 isolates each derived from a different patient.

**Figure 1.**
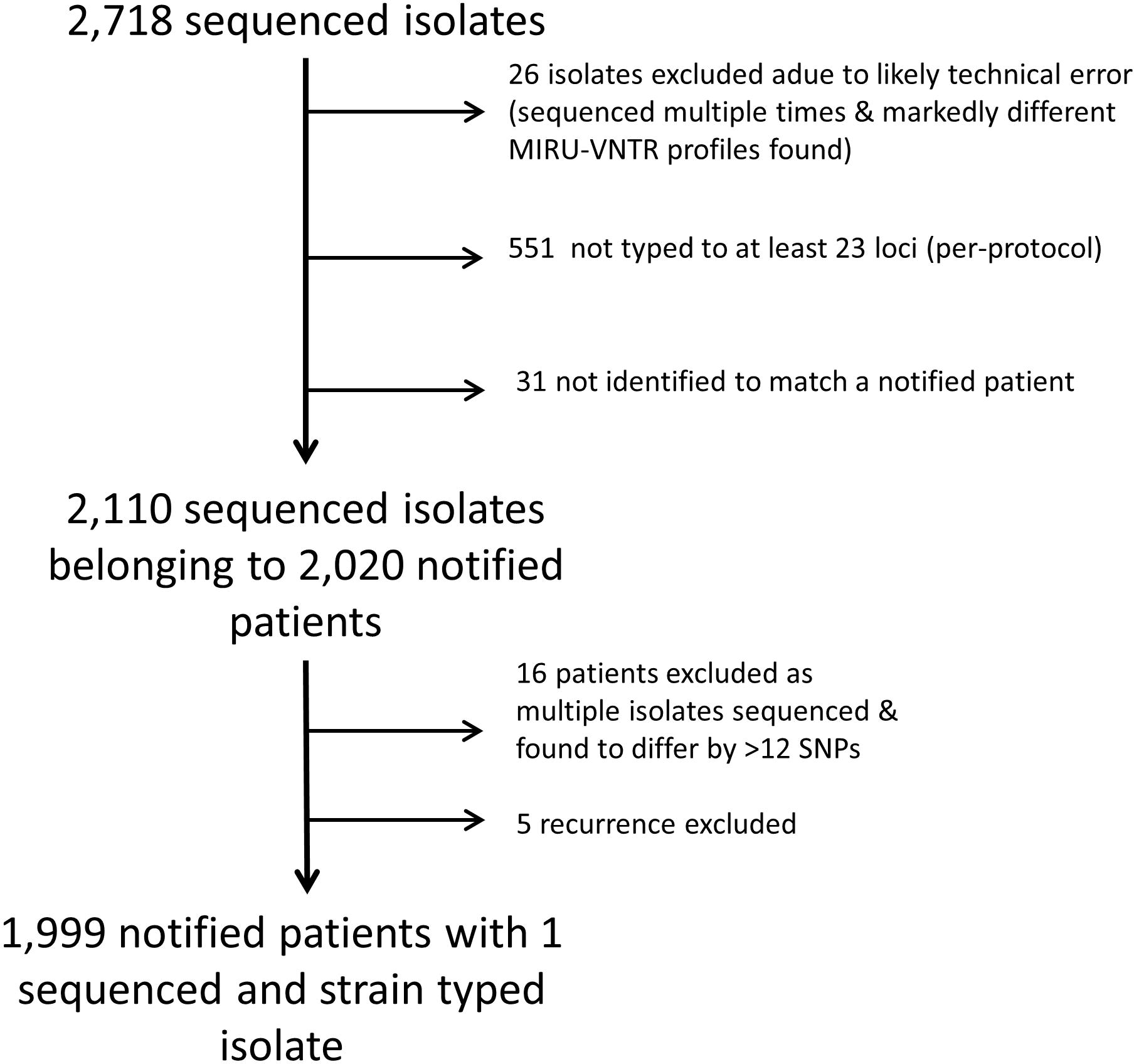
Flowchart showing the samples studied. Flowchart showing the samples studied.

There were more male than female patients (1176, 58%). 1155 (58%) were aged between 15-44 years old. 1325 patients (66%) were born outside the UK and 1437 (71%) were of non-White ethnicity (Table 2). *M. tuberculosis* lineage 4 (Euro-American) was the most commonly isolated lineage (n=954, 48%) with lineages 1, 2, and 3 also commonly represented (176 (9%), 137 (7%), 704 (35%) isolates respectively) (Table 2). *M. tuberculosis* lineage was associated with country of birth, with lineage 3 being most common in individuals born in India or Pakistan (Table 3).

**Table 2.** Details of Samples studied

| Category | Property | Number of samples |
| --- | --- | --- |
| Number of social risk factors<br>(homelessness, prison, alcohol use, drug use) | 0 | 1761 |
|  | 1 | 136 |
|  | 2 | 51 |
|  | 3 | 26 |
|  | 4 | 3 |
|  | Not available | 22 |
| Gender | Female | 801 |
|  | Male | 1176 |
|  | Not available | 22 |
| Age group | 0-14 | 45 |
|  | 15-44 | 1155 |
|  | 45-64 | 442 |
|  | 65+ | 335 |
|  | Not available | 22 |
| Year sample taken | 2007 | 1 |
|  | 2010 | 1 |
|  | 2011 | 5 |
|  | 2012 | 355 |
|  | 2013 | 584 |
|  | 2014 | 507 |
|  | 2015 | 524 |
|  | Not available | 22 |
| PHE Region of patient's residence | London | 6 |
|  | Midlands & East of England | 1721 |
|  | North of England | 243 |
|  | South of England | 3 |
|  | Not available | 26 |
| Self-declared ethnic group | Bangladeshi | 31 |
|  | Black-African | 267 |
|  | Black-Caribbean | 57 |
|  | Black-Other | 14 |
|  | Chinese | 29 |
|  | Indian | 564 |
|  | Mixed / Other | 143 |
|  | Pakistani | 332 |
|  | White | 508 |
|  | Not available | 54 |
| UK Born | Non-UK Born | 1325 |
|  | UK Born | 592 |
|  | Not available | 82 |

**Table 3.** Lineage of isolates studied

| Place of birth | Lineage |  |  |  |  | Total |
| --- | --- | --- | --- | --- | --- | --- |
|  | 1 | 2 | 3 | 4 | Other |  |
| UNITED KINGDOM | 19<br>(3.2%) | 33<br>(5.5%) | 136<br>(23%) | 391<br>(66%) | 13<br>(2.1%) | 592<br>(100%) |
| INDIA | 81<br>(18%) | 18<br>(4.0%) | 246<br>(55%) | 102<br>(23%) | 1<br>(0.2%) | 448<br>(100%) |
| PAKISTAN | 16<br>(6.3%) | 6<br>(2.3%) | 178<br>(71%) | 51<br>(20%) | 1<br>(0.4%) | 252<br>(100%) |
| SOMALIA | 8<br>(15%) | 2<br>(3.8%) | 24<br>(45%) | 18<br>(33%) | 1<br>(1.9%) | 53<br>(100%) |
| ZIMBABWE | 3<br>(6.0%) | 7<br>(14%) | 1<br>(2.0%) | 37<br>(76%) | 1<br>(2.0%) | 49<br>(100%) |
| ERITREA | 3<br>(6.5%) | 2<br>(4.3%) | 16<br>(35%) | 25<br>(54%) | 0<br>(0.0%) | 46<br>(100%) |
| POLAND | 0<br>(0.0%) | 1<br>(2.8%) | 1<br>(2.8%) | 34<br>(94%) | 0<br>(0.0%) | 36<br>(100%) |
| ROMANIA | 0<br>(0.0%) | 0<br>(0.0%) | 0<br>(0.0%) | 28<br>(100%) | 0<br>(0.0%) | 28<br>(100%) |
| LITHUANIA | 0<br>(0.0%) | 8<br>(33%) | 0<br>(0.0%) | 16<br>(66%) | 0<br>(0.0%) | 24<br>(100%) |
| Other | 36<br>(9.5%) | 58<br>(15%) | 66<br>(18%) | 208<br>(55%) | 9<br>(2.4%) | 377<br>(100%) |
| Not known | 10<br>(10%) | 2<br>(2.1%) | 36<br>(38%) | 44<br>(49%) | 2<br>(2.1%) | 94<br>(100%) |
| Total | 176 | 137 | 704 | 954 | 28 | 1999 |

### Epidemiological risk factors and the prediction of close relatedness

Using pairwise SNV distances within 5 SNVs between isolates to define genomic relatedness, we determined how various shared epidemiological data altered the odds of relatedness. Figure 2A shows estimated odds ratios of close genomic relatedness, in the presence, relative to the absence, of a series of risk factors. The proportion of paired isolates that are closely genomically related, given a particular risk factor, was also calculated. This represents the positive predictive value (PPV) of each risk factor. SNV thresholds other than 5 SNVs were analysed in sensitivity analyses (Web extra Fig. S1-S6), with similar results.

**Figure 2.**
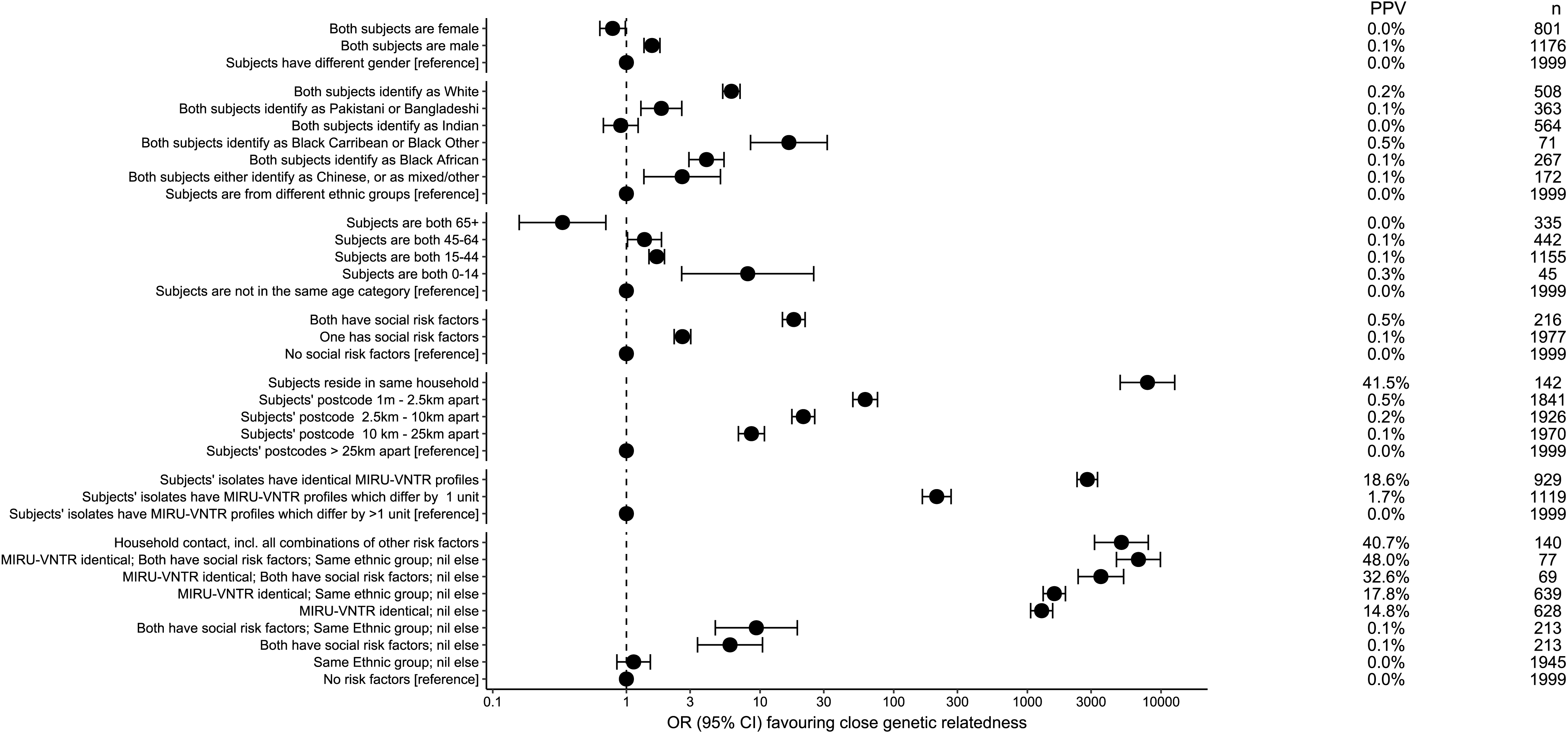
Relationship between MIRU-VNTR profile, epidemiological risk factors and genetic relatedness. The odds ratio predicting closely related isolates (defined by having five or fewer single nucleotide variants between them) associated with sharing a series of epidemiological properties. PPV denotes positive predictive values. n refers to the number of subjects having the property described. For example, there were 801 female subjects.

Predictably, residence at the same address was most strongly associated with close genomic relatedness (OR 8,000, 95% CI 5,000, 13,000). This corresponds to a PPV of 42%, indicating the majority of co-resident cases in this series were not closely genomically related, something discussed below. However, it was rare for two patients to share an address, with only 85 isolates derived from such settings. Other risk factors studied included sharing a self-identified ethnic group with another patient or being in a similar age bracket. Both were weakly associated with genomic relatedness (estimated odds ratios of 10 or less), with the highest risk of close genomic relatedness for an ethnic group seen for the smallest ethnic group studied (those identifying as Black Caribbean or Black Other; n=71; OR 16, 95% CI 8, 32). Similarly, there was a modest increase in the odds of close genomic relatedness where two isolates were from individuals with social risk factors (current or history of imprisonment, drug misuse, alcohol misuse or homelessness) (OR 9, 95% CI 4, 16). In all these cases however, the PPV was less than 1%.

### MIRU-VNTR profiles as predictors of close relatedness

Having identical MIRU-VNTR profiles conferred an odds ratio of close genomic relatedness of 2,800 (95% CI 2,200, 3,400) on paired isolates, compared with paired isolates with different MIRU-VNTR profiles, with an associated 18.6% PPV (Figure 2B). With 1 locus discordant, the corresponding odds ratio and PPV were much lower (OR 210, 95% CI 160,270; PPV 1•7%).

To understand how MIRU-VNTR profile and epidemiological data can complement each other in the identification of close relatedness, we assessed combinations of the presence of identical MIRU-VNTR profiles, social risk factors, and shared ethnicity, all factors which are significantly associated with close relatedness individually (Figure 2A). Excluding individuals who were resident at the same address, identical MIRU-VNTR profile was more predictive of close relatedness when shared risk factors were present, but for all the combinations studied the PPV remained low (15%, 18%, 33%, 48% with no shared risk factors, same ethnic group but no social risk factors, shared social risk factors but different ethnic group, and both shared ethnic group and social risk factors, respectively).

### SNV - MIRU-VNTR relationships vary by lineage

While MIRU-VNTR profiles predict close genetic relatedness (defined by SNVs) better than most social risk factors (Figure 2A), we observed that the PPV differs markedly by *M. tuberculosis* lineage (Fig. 3). For lineages 1, 2, 3 and 4, which together account for 1,977/1,999 (99%) of the isolates studied, we compared pairwise comparisons within each lineage by MIRU-VNTR similarity (Fig. 4). For lineages 1 and 4, pairwise SNV distances increased over the range 0 to 8 MIRU-VNTR unit differences, until at higher MIRU-VNTR distances the pairwise distances approximated the within-lineage median pairwise SNV distance (Fig. 4). For lineages 2 and 3 the median was reached by 3 MIRU-VNTR differences. Overall there was less variation between paired isolates within lineages 2 and 3 (median pairwise distances 205 and 334, respectively) compared to paired isolates within lineages 1 and 4 (median pairwise distances 840, and 685). However, for paired isolates differing by between zero and 4 MIRU-VNTR loci, the least variation was seen within lineage 4.

**Figure 3.**
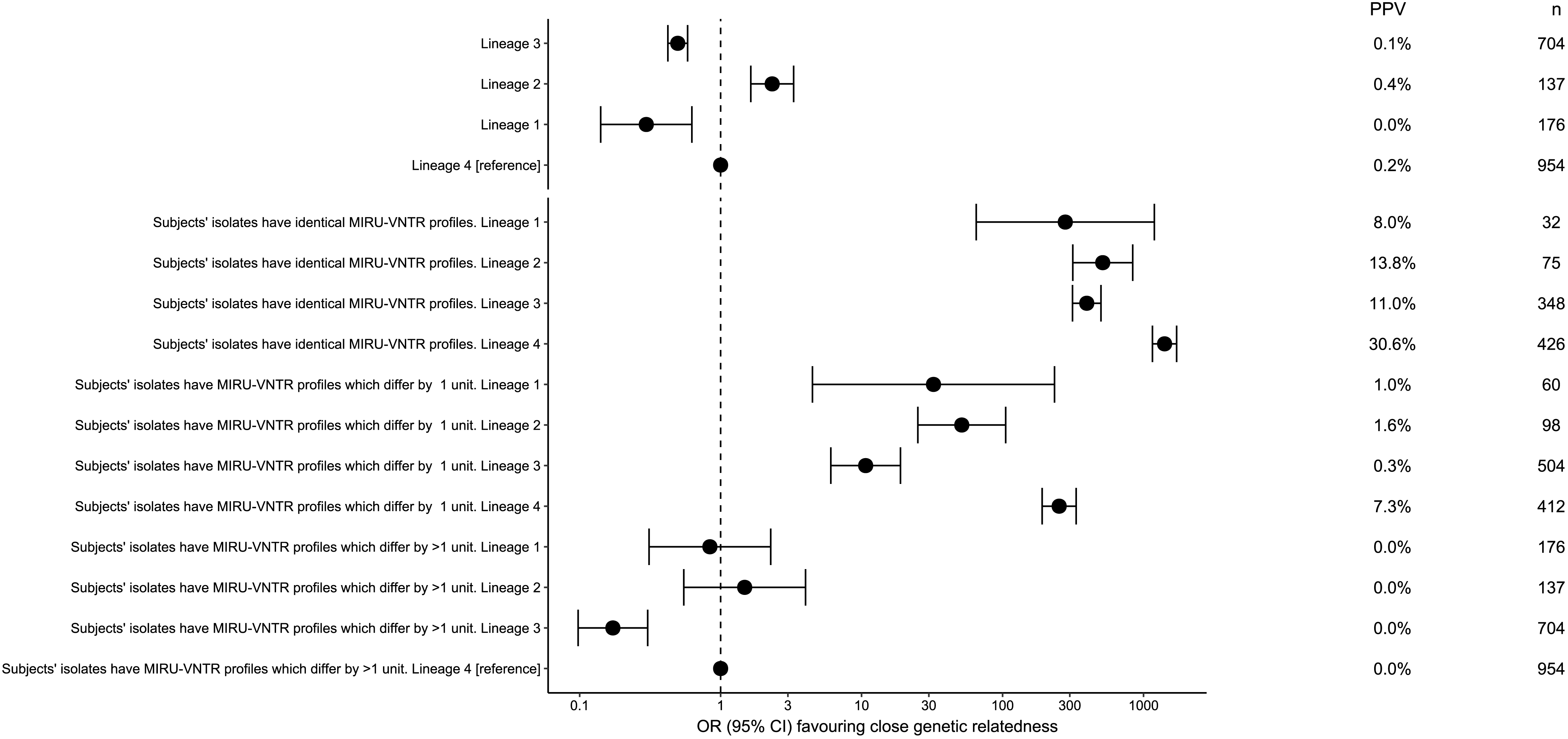
Association between lineage, close genetic relatedness and MIRU-VNTR profile. The odds ratio predicting closely related isolates (defined by having five or fewer single nucleotide variants between them) associated with sharing a particular lineage (relative to lineage 4), or having identical or similar MIRU-VNTR profiles, stratified by lineage. PPV denotes positive predictive values. n refers to the number of subjects having the property described. For example, there were 954 subjects of lineage 4.

To quantify how the relationship between MIRU-VNTR and SNVs differed by lineage, we modelled SNV distances between paired isolates, assuming a linear relationship with MIRU-VNTR profile distances over the range of 0-3 MIRU-VNTR unit differences (Figure 4, red dots show fitted medians, and Supplementary Data 2). For lineage 4 isolates, among pairs with identical MIRU-VNTR profiles, there was a median of 10 ± 0•4 SNV (median ± standard error). For paired isolates with identical MIRU-VNTR profiles in lineages 1, 2, and 3, SNV distances were 122 ± 21, 159 ± 3, and 82 ± 3 (median ± standard error), respectively. According to current estimates of *M. tuberculosis* clock rates, these correspond to about 250, 300, and 150 years of evolution, respectively, compared to about 20 years for lineage 4^19^.

**Figure 4.**
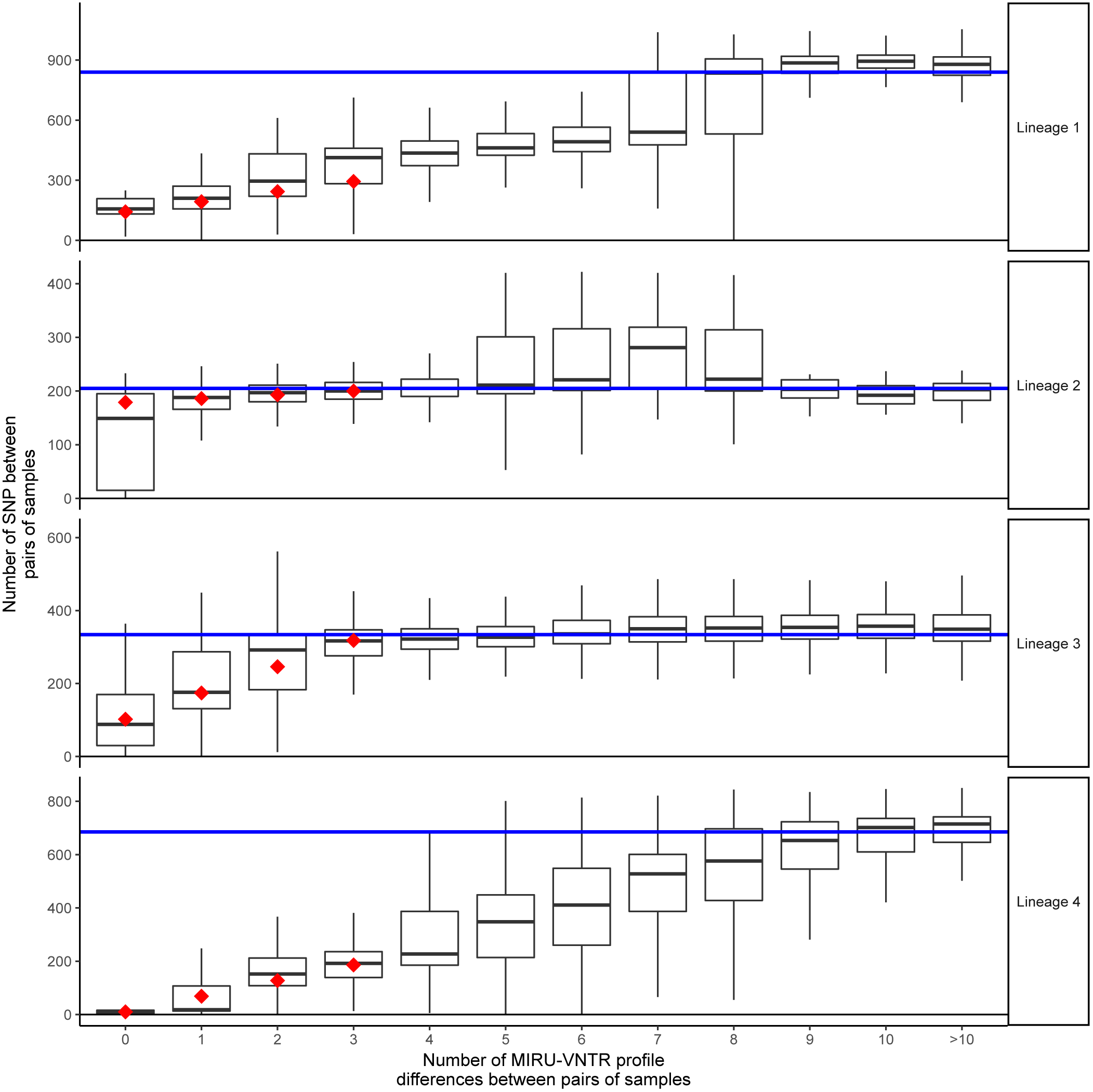
The relationship between lineage, MIRU-VNTR profile variation and SNV variation. The relationship between MIRU-VNTR profile variation and SNV variation, stratified by lineage. The x-axis shown the number of MIRU-VNTR repeats differing between pairs of isolates. For example, if a sample had a MIRU-VNTR profile of 121, and another 111, locus #2 has reduced in repeat number by one, which counts as a 1 MIRU-VNTR profile repeat number change. The y-axis shows the median number of SNV in each of a large number of pairs examined. The blue line reflects the median pairwise distance within all sampled isolates of each lineage. Red dots are fitted median values from a multivariable model fitted to MIRU-VNTR profile differences between 0 and 3.

For each MIRU-VNTR unit difference in lineage 4, there was a median increase of 59 ± 0•6 SNV. For lineage 1, a similar increase in SNV with increasing MIRU-VNTR differences was observed to that in lineage 4 (het*. p* = 0•32), whereas for lineages 2 and 3 the relationship was very different from lineage 4 (het. *p* < 10^-20^ for both comparisons). Indeed, for paired isolates in lineage 2, SNVs were only weakly associated with MIRU-VNTR distance. Thus, in the population studied, the performance of MIRU-VNTR profiles in defining evolutionarily related groups differed between lineage 4 (Euro-American) isolates, and lineages 1, 2 and 3.

## Discussion

In this prospective study of a cosmopolitan population in the English Midlands, we have quantified how well recent transmission, as defined a 5 SNV threshold, is predicted by shared epidemiological risk factors, by MIRU-VNTR typing, or by a combination of both.^19^ We have also demonstrated how lineage strongly affects the performance of MIRU-VNTR-based predictions.

Overall, the PPV for recent transmission for any two isolates with an identical MIRU-VNTR type was only 18.6%. Excluding cases resident at the same address, the PPV varied from as low as 14.8% to 48.0% if shared risk factors were present alongside identical MIRU-VNTR profiles (Figure 2). However, PPVs for shared MIRU-VNTR profiles differed significantly by lineage, with the strongest associations seen in lineage 4 (European-American), which was also most frequently observed lineage in the Midlands. The number of patient-to-patient links that need to be investigated to find a single case of recent transmission between non-co-resident individuals with shared MIRU-VNTR types is thus between two and seven, depending on the presence of shared social risk factors.

These data demonstrate that the previous routine practice of grouping samples based on MIRU-VNTR identity, or on a combination of MIRU-VNTR identity and shared epidemiological risk factors, generates highly heterogeneous results, and is likely to contribute to the low cost-effectiveness of MIRU-VNTR typing.^7^ Importantly, our data also demonstrate how lineage markedly affects the PPV of MIRU-VNTR links, with the best results seen for lineage 4. To our knowledge, lineage has not been routinely taken into consideration when matching isolates by MIRU-VNTR for surveillance reasons.

One possible explanation for why SNV distances between paired isolates sharing a MIRU-VNTR profile within lineages 1, 2 and 3 were greater than for lineage 4 is that the Indo-Oceanic, East-Asian (including Beijing) and East-African Indian lineages are more endemic to countries other than the UK, and that patients diagnosed with these tuberculosis lineages in the UK were infected overseas. Were this the case, closely genomically related strains would be less likely to be found in England. For example, lineage 3 isolates were most common in individuals born in India and Pakistan, relative to other individuals, supporting this hypothesis (Table 3). A second possible explanation is that the rate of diversification of MIRU-VNTR types relative to SNVs differs between major lineages. Thirdly, MIRU-VNTR variation can result in the same profile via different evolutionary routes (homoplasy)^20^, a phenomenon which could also explain the rather flat relationship observed between MIRU-VNTR distance and SNV distance seen in lineages 2 and 3. Whatever the mechanism(s) operating, our data implies that the lineages, and their epidemiology, may influence the wide variation in the proportion of TB cases clustering using MIRU-VNTR profiling reported in different settings.^9,21^

It was surprising to us that among individuals resident at the same address, only 42% of these pairs were closely genomically linked. One explanation for this relatively low proportion is that some patients from highly endemic countries are likely to co-habit with others from highly endemic countries, potentially increasing the chances of non-clustered isolates, originating from separate exposures, being linked to the same address. Another scenario that could lead to a similar effect would be UK born patients with multiple social risk factors sharing hostels. In both settings, co-resident individuals with TB would be expected to have an increased risk of having acquired their infection from individuals in high prevalence populations with whom they have been in contact outside the residential setting.

One limitation to this study is that is that the results cannot necessarily be generalised to other settings with different patterns of transmission, rates of disease, patterns of immigration, and relative prevalence of different lineages. However, the region studied was large and included a mixture of incidence areas, and both urban and rural settings. Another potential limitation is that we cannot be sure that risk factor data was recorded in a fully sensitive manner. Under-ascertainment of risk factor data would reduce the apparent contribution of risk factor data to identifying close genetic neighbours. However, even in the population in which we found in which MIRU-VNTR profiling works best (lineage 4 infections), and in subjects for whom shared risk factors were recorded, the combination of MIRU-VNTR identity and shared risk factors only detects about one in two closely related isolate pairs.

In summary, these data help quantify the limitations of MIRU-VNTR typing for tuberculosis transmission surveillance and control. With routine diagnostic services beginning to transition to WGS technology in multiple high incidence countries, as England already has, our data indicates one can expect to see a reduction in the number of potential links requiring epidemiological investigation by a factor of about five. WGS thus stands a much greater chance of contributing to a cost effective control program than MIRU-VNTR typing in low-burden, cosmopolitan settings such as ours.

